# Image Scanning Microscopy with Single-Photon Detector Array

**DOI:** 10.1101/335596

**Authors:** Marco Castello, Giorgio Tortarolo, Mauro Buttafava, Takahiro Deguchi, Federica Villa, Sami Koho, Paolo Bianchini, Colin J. R. Sheppard, Alberto Diaspro, Alberto Tosi, Giuseppe Vicidomini

## Abstract

**Image scanning microscopy (ISM) improves the spatial resolution of conventional confocal laser-scanning microscopy (CLSM), but current implementations reduce versatility and restrict its combination with fluorescence spectroscopy techniques, such as fluorescence lifetime. Here, we describe a natural design of ISM based on a fast single-photon detector array, which allows straightforward upgrade of an existing confocal microscope, without compromising any of its functionalities. In contrast to all-optical ISM implementations, our approach provides access to the raw scanned images, opening the way to adaptive reconstruction methods, capable of considering different imaging conditions and distortions. We demonstrate its utility in the context of fluorescence lifetime, deep, multicolor and live-cell imaging. This implementation will pave the way for a transparent and massive transition from conventional CLSM to ISM**.

**confocal microscopy | time-resolved spectroscopy | image scanning microscopy | single-photon detector array**

---

Fluorescence confocal laser-scanning microscopy (CLSM) is an indispensable tool for biomedical research by virtue of its versatility, its optical sectioning capability and its harmonization with spectroscopic assays. In CLSM, an excitation laser beam is focused to a diffraction-limited volume on the sample. The fluorescence from this volume is registered by a single-point detector, after being filtered by a pinhole (typically 1 Airy unit (A.U.) in radius) to reject the out-of-focus light. Finally, this detection volume is raster scanned across the sample to form the image. By recording the signal as a function of wavelength, polarization or/and time, the structural information provided by imaging can be correlated with different spectroscopic parameters, such as the fluorescence lifetime, to decipher structural and functional relations (1). Spatial resolution is another important asset of confocal microscopy: by closing the optical pinhole, the detection volume - or point-spread-function (PSF) - can <u>b</u>e shrunk down, up-to a limited improvement factor of 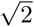 with respect to the regular pinhole-less widefield PSF. However, the shrinking of the PSF does not always translate into a practical resolution enhancement, since closing the pinhole also causes a heavy signal loss, and thus a reduction of the signal-to-noise ratio (SNR) of the image. More rigorously, CLSM theoretically doubles the imaging cut-off frequency of conventional microscopy, but the magnitudes of these extra-frequencies are usually lower than the noise level (2).

Image scanning microscopy (ISM), theoretically propose<u>d</u> almost thirty years ago (3), makes it possible to attain the 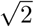 resolution enhancement predicted for CLSM, i.e., the magnitudes of the frequencies within the cut-off are effectively increased above the noise level (2). In short, in ISM the single point-detector of a confocal microscope is replaced with a detector array – and the pinhole is opened to ∼ 1 A.U. to collect most of fluorescent light, while maintaining optical sectioning (4). Hence, the detector generates a series of “confocal” scanned images of the same sample, but with different information content, and shifted with respect to each other according to the relative positions of the detector elements. These images are successively recombined together (through pixel-reassignment in the original ISM idea (3, 4) or equivalently through Fourier-based or deconvolution-based image reconstruction in some later works (5–7)) to form a final image with a PSF equivalent to - or even smaller than - the ideal (infinitely small pinhole) confocal microscope, but without discarding fluorescence photons, and thus without decreasing the SNR.

In the first implementation of ISM a conventional camera was used as the detector array (5), but its the frame-rate (read-out bandwidth) severely limited the imaging speed. Later, this limitation has been overcome by the so-called all-optical ISM implementations (8–11) - sometimes combining ISM with multi-spot excitation architectures (12). The pixel-reassignment method allows obtaining an enhanced-resolution image by adding together all the scanned images, after shifting each one of them by a vector depicting the position of the corresponding detector element, properly scaled by the pixel-reassignment factor. In all-optical implementations the shift operation is obtained by upscaling the position of the scanned images within the final image by the pixel-reassignment factor, and the superimposition is obtained by taking advantage of the integration of the camera during the raster scanning. Since the upscaling is obtained by using a second synchronized scanners (8, 10), or passing twice the emission beam across the same scanner (9), such implementations substantially modify the architecture of conventional CLSM, reducing its versatility. A simplified all-optical ISM architecture has been recently proposed (11); however, it can not implement a pinhole, and thus optical sectioning is obtained only by using non-linear contrast mechanisms such as two-photon-excitation. Furthermore, in all-optical ISM implementations the pixel-reassignment factor needs to be decided *a-priori*, i.e., before imaging. In this context, it is important to highlight that: (i) for some all-optical implementations (9, 11), the pixel-reassignment factor is fixed by the configuration of the optical architectures, thus its modification is not straightforward; (ii) the pixel-reassignment factor is equal to two only in the ideal case of identical excitation and emission PSFs. Any changes in the shape of the PSFs, caused by system distortions or the sample, change the reassignment factors, and thus their effective estimation *a-priori* is not feasible. Finally, all-optical implementations do not give access to the raw scanned images, and, since they use camera devices, combination with spectroscopic techniques, such as fluorescence lifetime imaging is not straightforward. Another approach to solve the scan speed limitation of the first ISM implementation is to use a pixelated point detector – a “camera” with a reduced number of elements – which allows improving the read-out-bandwidth – and to implement the original ISM principle with a minimal change of the conventional CLSM architecture. The first attempt in this direction was the AiryScan system (13), in which to mimic a detector array, a hexagonal-shaped bundle of optical fibers is connected to a one-dimensional array of GaAsP photo-multiplier-tubes (PMT) working in linear (or analog) mode. However, in recent years, microelectronic detectors, like single-photon avalanche diodes (SPADs), are becoming increasingly popular in the field of time-resolved spectroscopy, thanks to their photon-counting nature (14, 15).

With performance comparable (or even superior) to PMTs in terms of detection efficiency, noise and temporal response, they are favored by the great scalability, robustness, flexibility and reliability offered by the microelectronic fabrication technology. Compared to their vacuum based counterparts, multi-pixel microelectronic single-photon detectors also contribute to reduce cost, overall size and system complexity. All these properties currently make the SPAD array one of the best technology to develop a real photon-counting bidimension detector array for fluorescence microscopy.

Here, we demonstrate a fully automatic, time-resolved ISM implementation based on a novel, fast and robust SPAD array (5 × 5 elements), specifically designed for this application (Supplementary Fig. S1, Supplementary Note 1). By a simple modification of the microscope detection arm (Fig. 1a), this SPAD array allows the transformation of any existing confocal system into an image scanning microscope, while preserving all its functionalities, such as optical sectioning, multi-color imaging, and fast imaging. The availability of all the raw scanned images allows for a self-calibrating pixel-reassignment method, able to reconstruct the high-resolution image also in case of unknown and/or variable pixel-reassignment factor. Furthermore, the single-photon detection ability (< 200 ps timing jitter) of each element of the SPAD array allows the combination of ISM with fluorescence lifetime (*via* the time-correlated-single-photon-counting method, TCSPC (15)).

**Fig. 1.**
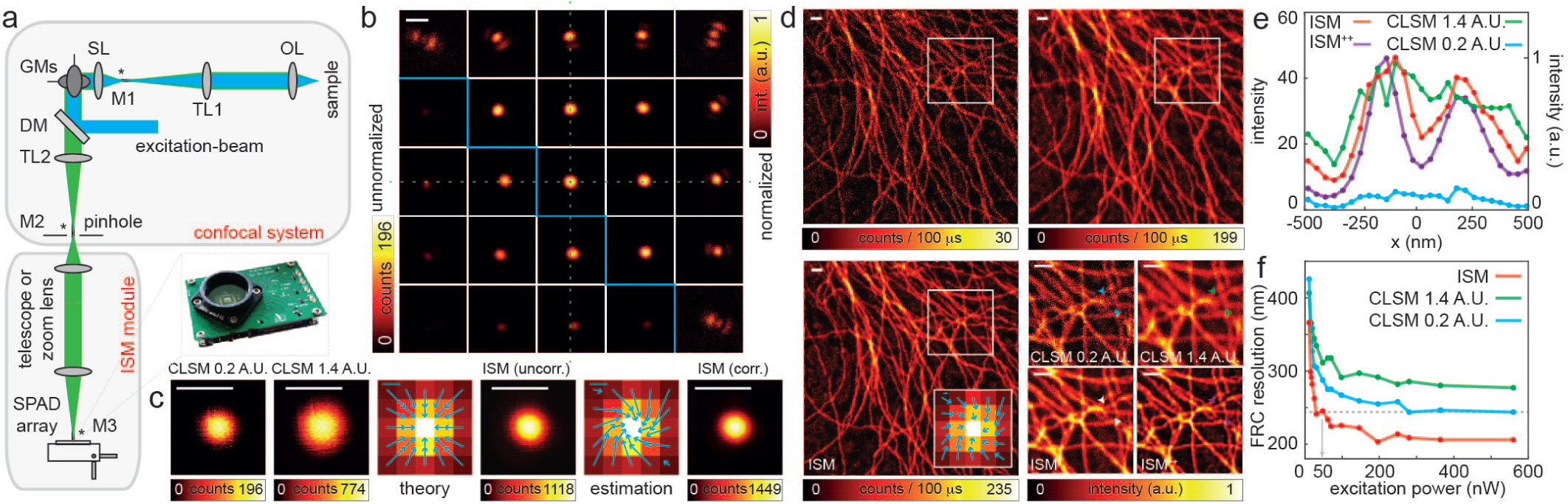
Image scanning microscopy with SPAD array. (a) Schematic of the image scanning microscope. Excitation light (blue) passes the dichroic mirror (DM) and is deflected by the galvanometer mirrors (GMs). The pivot point of the scanner is projected by the scan and tube lenses (SL, TL1) into the back aperture of the objective lens (OL). Fluorescence (green) is collected by the objective lens, de-scanned by the GMs, filtered by the DM and projected by a second tube lens (TL2) into the pinhole plane. A telescope or a zoom lens is used to image the pinhole plane into the SPAD array and add an extra magnification (from M2 to M3) to match the physical size of the detector to 1.4 A.U. The asterisks denote the planes conjugated. The picture shows the printed circuit board hosting the detector. (b) Matrix representing the scanned reflection images of a single isolated gold bead (80 nm). Each scanned image of the top-right corner and the bottom-left corner are normalized to its maximum intensity and to the maximum intensity of the central scanned image, respectively. Horizontal and vertical dashed-green lines guide the eye. Scale bar: 500 nm. (c) Side-by-side comparison of the PSF of the “ideal” confocal (0.2 A.U. pinhole), “open” confocal (1.4 A.U. pinhole), uncorrected ISM (i.e., pixel-reassignment with the theoretical shift-vectors) and corrected pixel-reassignment ISM (i.e., pixel-reassignment with the estimated shift-vectors) obtained from (c). Scale bars: 500 nm. The fingerprint maps of (c), superimposed with the theoretical and estimated shift-vectors projected in the image plane, are also shown. Scale bars: 50 nm. (d) Side-by-side comparison of “ideal” confocal, “open” confocal, and ISM images of tubulin filaments stained with Abberior STAR red. Pixel-dwell time: 50 *µ*s. Pixel-size: 37.6 nm. Image format: 400 × 400 pixels. Excitation power *P*_*exc*_ = 280 nW. Scale bars: 1 *µ*m. Inset shows the fingerprint map and the estimated shift-vectors. Scale bar: 100 nm. Magnified views of the dashed box area are also reported, together with the ISM image obtained using multi-image deconvolution (ISM^++^, 5 iterations). Excitation power *P*_*exc*_ = 280 nW. Scale bars: 1 *µ*m. (e) Line intensity profiles across two branching tubular filaments at the position of the arrowheads for the different imaging modalities. (f) FRC-based resolution as function of the excitation intensity beam for the different imaging modalities.

We first integrated our SPAD array into a custom confocal laser scanning microscope (Supplementary Fig. S2); its pinhole is completely opened and its single-point detector is substituted by a telescope and a SPAD array. The telescope conjugates the pinhole with the SPAD array and provides a magnification such that the size of the SPAD array projected onto the sample plane is ∼ 1 A.U. Practically, the detector mimics a 1 A.U. pinhole, allowing to maintain the optical sectioning ability of the microscope also in the case of single-photon excitation. As a first demonstration of ISM with the SPAD array we imaged a single isolated sub-diffraction (80 nm in diameter) gold bead in reflection mode, giving us access to the PSF of the proposed implementation. Given the 25 raw scanned images (Fig. 1b) the high-resolution image is obtained by using the pixel-reassignment method based on the prior knowledge of the system (Fig. 1c); (i) each scanned image is shifted by the vector representing the distance between the associated SPAD array element and the “central” element (which amplitude is proportional to the pixel-pitch of the array), divided by the magnification on the detector, the pixel size of the image and the pixel-reassignment factor, which is equal to 2 in case of reflection (illumination and detection PSF are identical); (ii) all shifted images are added together. The PSF of the “ideal” confocal (0.2 A.U., i.e., scanned image from the central element) is reduced in size with respect to the “open” confocal (1.4 A.U., i.e., all scanned images are added up without any shift) counterpart - from 281 nm to 213 nm (FWHM), but the peak intensity substantially reduces. On the contrary, the PSF of the ISM system reduces in size (217 nm, FWHM) and improves in peak intensity, both with respect to the “ideal” (∼ 5.7 fold) and the “open” (∼ 1.4 fold) confocal (Supplementary Fig. S3). This latter enhancement is known as the super-brightness effect (4). To highlight the importance to have access to the scanned images - which is precluded in all-optical implementations -, we derived a method based on phase-correlation (6, 16), which estimates the shift-vectors for the pixel reassignment directly from the scanned images (Fig. 1c, Supplementary Note 2), making the prior knowledge of the pixel-reassignment factor, and more in general the need of theoretical shift-vectors, obsolete. By using this automatic and self-calibrating approach, the PSF of the image scanning microscope further reduces its size (to 193 nm) and improves its peak intensity (∼ 1.8 and ∼ 7.4 fold, with respect to “open” and “ideal” confocal, respectively). The same method is also useful to automatically compensate for the fluorescent Stoke-shift of fluorescence microscopy. Contrary to reflection microscopy, illumination/excitation and emission PSFs change, thus also in the case of optimal imaging conditions the reassignment factor is not exactly 2, but depends on the Stokes-shift between the excitation and emission wavelength (17).

Because of the ancillary SNR boost and the PSF’s size reduction, the resolution of the ISM image is clearly enhanced as compared with both “ideal” and “open” pinhole confocal images (Fig. 1d, Supplementary Fig. S4). Since the ISM image results from the sum of the scanned images, it is still expressed in photon counts and is linear with the fluorophore’s concentration, which make this approach fully compatible with a quantitative analysis (18). We quantify the resolution enhancement by plotting the line intensity profiles across tiny and close-packed filaments (Fig. 1e) and *via* the Fourier-ring correlation (FRC) method (19). The FRC analysis is able to consider both the PSF’s size reduction and, more important in this context, the SNR enhancement. For example, the FRC-based resolution as a function of the excitation beam power (Fig. 1f, Supplementary Fig. S5) shows the ability of ISM to achieve the same resolution of “ideal” CLMS at one tenth of the illumination intensity a great benefit for live cell imaging (Supplementary Fig. S6, Supplementary Video 1-4). Given the raw scanned images and the shift-vectors, the high-resolution image can be also reconstructed *via* multi-image deconvolution (20, 21) (Supplementary Note 4), which offers a further enhancement of SNR and effective resolution 1(d). The access to the raw data also allows the derivation of important system information, such as the effective magnification (Supplementary Note 3) and the optical status. Regarding the microscope status, the so-called fingerprint map, i.e., the map representing the total signal of each scanned image, encodes information about both the PSF (Fig. 1d, Supplementary Note 3) and the detector misalignment.

To demonstrate that the proposed ISM implementation is compatible with time-resolved spectroscopy, in particular with techniques that require detecting single-photons with low-time jitter such as fluorescence lifetime (FL), we registered the signal from the elements of the SPAD array with a multi-channel TCSPC card able to synchronize the registration of each photon with the pulses of the excitation source and the beam scanning system (time-tag modality). In the context of FL image scanning microscopy (FLISM) (Fig. 2), the time-tag data are analyzed to produce a series of three-dimensional scanned images, where the third dimension (temporal-bin) is the photon-arrival time histogram (Fig. 2a). Given this series of scanned images, the pixel-reassignment procedure is applied for each temporal bin and the resulting three-dimension ISM dataset is processed using a conventional FL fitting approach to generate the FL map. The final result is a FL map/image with higher spatial resolution and higher lifetime precision with respect to the confocal counterpart (Fig. 2b) thank to the PSF shrinking and the super-brightness effect.

**Fig. 2.**
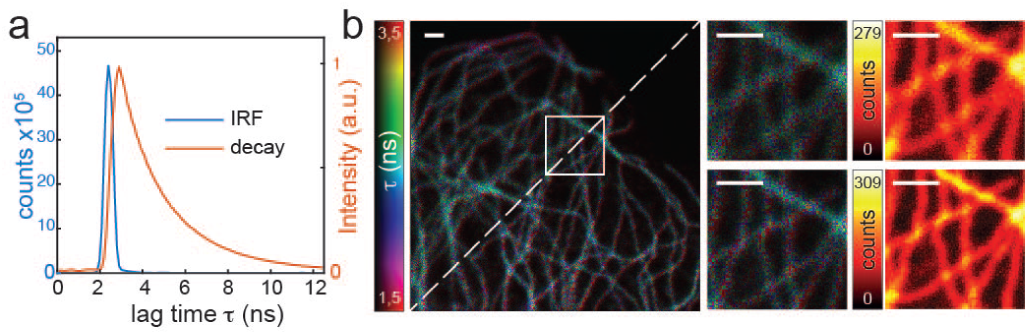
Fluorescence lifetime image scanning microscopy. (a) The impulse response function (IRF) of the image scanning microscope, measured by using the reflection from a gold-bead, and the photon-arrival-time histogram for tubulin labeled with Abberior STAR red. (b) Side-by-side comparison of the “open” confocal FLIM (top-left) and the FLISM (bottom-right) images of tubulin labeled with Abberior STAR red. Magnified views of the white box for the FLIM and FLISM images. The respective intensity images are also reported. Pixel-dwell time: 100 *µ*s. Pixel-size: 30 nm. Image format: 500 × 500 pixels. Excitation power *P*_*exc*_ = 500 nW. Scale bars: 1 *µ*m.

Next to demonstrate that our ISM implementation can effectively pave the way for a non-invasive and straightforward transition from conventional CLSM to ISM, we integrated the SPAD array into a commercial confocal microscope, by taking advantage of the typical extra detection ports of modern confocal scanning heads. Thanks to a zoom lens system instead of a fixed telescope system - we generated a image plane – conjugated with the pinhole – outside the scanning head of the confocal microscope where we located our SPAD array. The use of a motorized zoom lens allows fast changing the magnification (M3) on the detector, which is mandatory when switching between different objective lenses, and/or useful when changing the detection aperture size as often happens in commercial microscopes. As an example, we used a 20 × water (Fig. 3a) and 10 × glycerol (Fig. 3b) objective lenses to image (down to 0.9 mm and 1.8 mm for 20 × and 10 ×, respectively) E-YFP expressing neurons in the cleared whole brain of a mouse. While the 1 A.U. projected size of the detector array allows maintaining the optical sectioning capability of the architecture, the pixel-reassignment provides terrific SNR enhancement giving effective access to the resolution expected from “ideal” CLSM. Thanks to the raw scanned images the shift-vectors for the pixel-reassignment can be calculated section-by-section, thus our ISM architecture compensates for different optical aberrations and misalignments that can occur during three-dimensional deep imaging (Supplementary Fig. S8). Another case in which the access to raw scanned images is important is “simultaneous” multicolor ISM (Fig. 3 c). Due to the different Stokes shifts of the fluorophores and potential differences in the system condition for the two excitation beams, the pixel-reassignment factor changes - more generally the shift-vectors change - which imposes a sequential imaging strategy. Since reconstruction is performed *a-posteriori*, pulse-by-pulse, pixel-by-pixel or line-by-line multi-color strategies can be implemented in the proposed ISM architecture. Notably, the possibility to estimate the shift-vectors directly from the scanned images is fundamental when combining ISM with stimulated-emission-depletion (STED) microscopy (22): in this case, the “excitation” and emission PSFs are substantially different and the shape of the former is influenced by many imaging parameters difficult to know (19).

**Fig. 3.**
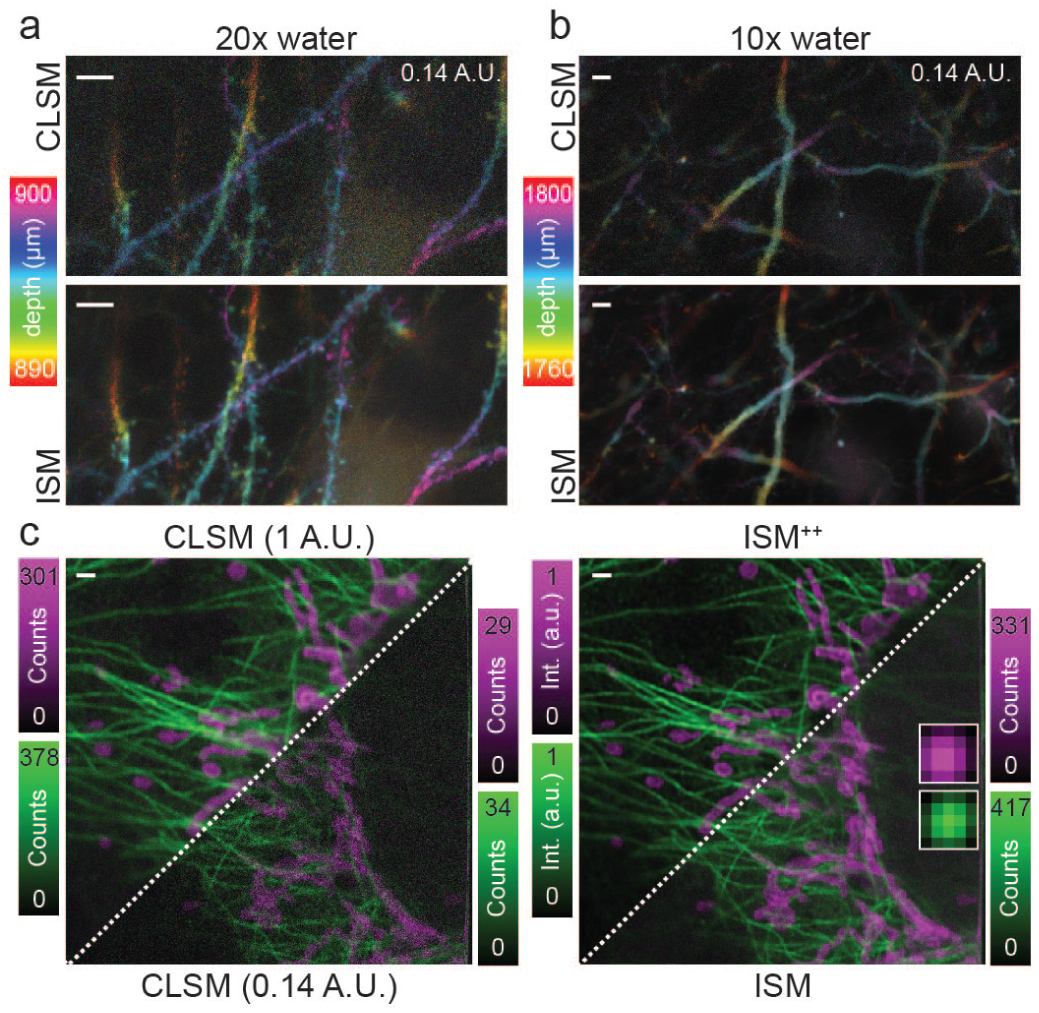
Image scanning microscopy with SPAD array on commercial confocal system. (a,b) Deep ISM imaging of neuronal processes in a whole cleared brain of Thy1-eYFP-H transgenic mouse with 20× (a) and 10× (b) water immersion objective lens. Side-by-side comparison of “ideal” confocal (top) and ISM (bottom) images of the maximum intensity projection - color coded by depth. Scale bars: 5 *µ*m. (c) Multi-color ISM of Alexa 488 for microtubules (green) and Alexa 568 for mitochondria (magenta). Side-by-side comparison of “open” confocal, “ideal” confocal, deconvolved ISM^++^ and pixel-reassignment ISM. Insets show the different fingerprint matrices for the two colors. Pixel dwell-time: 30 *µ*s. Scale bars: 1 *µ*m.

We have demonstrated a straightforward and non-invasive implementation of ISM based on a novel SPAD array. This implementation supports any objective lens and maintains optical sectioning, “simultaneous” multi-color imaging, live-cell imaging, and deep imaging. Thanks to the access to the raw scanned images, our method can extract system parameters, which is fundamental to obtain high-resolution images in the case of system distortions and unknown Stokes shift of the fluorophores. More in general, the analysis of the raw data provides important information regarding the microscope. For example, the fingerprint map encodes PSF information and this may trigger novel methods for correcting optical aberration *via* adaptive optics (23) and/or novel user-friendly deconvolution algorithm where no-prior information about the PSF are needed. In contrast to the all-optical implementation, our fully-automatic reconstruction approach does not allow true real-time imaging. However, the low computational complexity of the pixel-reassignment method concedes the high-resolution image immediately after the scanning. Otherwise, by using pre-loaded shift-vectors, for example obtained from previous scanned images, the high-resolution image can be reconstructed in real-time (pixel-by-pixel). The photon-counting detection ability of the SPAD array allows the integration of ISM with different time-resolved assays, and we have demonstrated the combination with FL imaging. However, the architecture can also be used for fluorescence correlation spectroscopy, other fluctuation correlation methods (24) and anti-bunching analysis (25). Another important temporal characteristic of the SPAD array is the absence of frame-rate (event-driven or asynchronous camera), namely, each element of the detector array is fully independent and fires a digital signal for each photon collected with a maximum read-out bandwidth of 40 MHz (25 ns hold-off time after registering a photon). These characteristics make the SPAD array compatible with smart illumination scheme (26) and with fast beam scanning systems, such as resonant scanners (Supplementary Fig. S9). Regarding the resonant scanner, it is normally combined with PMTs, due to their higher dynamic range as compared with single-photon detectors. However, since the photons emitted by the excitation volume are spread across all the elements of the SPAD array, the effective dynamic range of the SPAD array is significantly higher than those of a single SPAD element.

In conclusion, we believe that the SPAD array can became the standard detector for a versatile and multi-parameter image scanning microscope, capable of replacing any conventional (confocal) point scanning microscope.

## ACKNOWLEDGEMENTS

This work has been partially supported (G.T. and G.V.) from the Compagnia di San Paolo (codice ROL 20641). We thanks Ryu Nakamura (Nikon Corporation, Japan) for supporting on confocal A1R measurements, Nikon Corporation for sharing useful technical information about confocal A1R and for dual-color (mitocondria and tubulin) sample preparation. We thanks Dr. Ryosuke Kawakami, Dr. Kohei Otomo, and Prof. Tomomi Nemoto (Research Institute for Electronic Science, Hokkaido University, Japan) for advises in whole brain imaging. We thanks Dr. Michele Oneto (Nikon Imaging Center, IIT, Italy) for live-cell imaging support.

## AUTHOR CONTRIBUTIONS

M.C and G.V. conceived the idea. A.T and G.V. supervised the project, with support from C.J.R.S, P.B. and A.D. M.C, M.B., F.V., A.T. and G.V. designed the SPAD array. F.V. and A.T. realized and characterized the SPAD array. M.C., G.T. and G.V. designed and implemented the custom ISM system. T.D., P.B. and G.V. integrated the ISM method on the commercial microscope. M.C., G.T and G.V. designed and developed the controlling architecture. M.C., G.T. S.K and G.V. developed the image processing and analysis software. M.C., G.T., T.D., P.B., and G.V. performed the ISM experiments and analyzed the data with support from all authors. M.C., G.T., and G.V. wrote the manuscript with input from all authors.

## ONLINE METHODS

**Custom microscope**. We implemented the custom-based ISM system (Supplementary Fig. S2) by modifying an existing point-scanning microscope previously described in (1). Briefly, the excitation beam was provided by a triggerable pulsed diode laser (LDH-D-C-640, Picoquant) emitting at 640 nm; we controlled its power thanks to an acoustic optical modulator (AOM, MT80-A1-VIS, AA opto-electronic). After being reflected by a dichroic mirror (H643LPXR, AHF Analysentechnik), the excitation beam was deflected by two galvanometric scanning mirrors (6215HM40B, CT Cambridge Technology) and directed toward the objective lens (CFI Plan Apo VC 60 ×, 1.4 NA, Oil, Nikon) by the same set of scan and tube lenses used in a commercial scanning microscope (Confocal C2, Nikon). The fluorescence light was collected by the same objective lens, descanned, and passed through the dichroic mirror as well as through a fluorescence band pass filter (685-70, AHF Analysentechnik). A 300 mm aspheric lens (Thorlabs) focuses the fluorescence light into the pinhole plane generating a conjugated image plane with a theorethical magnification of 300 ×. For ISM measurements the pinhole was maintained completely open. A telescope system built using two aspheric lenses of 100 mm and 150 mm focal length (Thorlabs) conjugates the SPAD array with the pinhole and provides an extra magnification factor. The final magnification on the SPAD array plane is 450 ×, thus the size of the SPAD array projected on the specimen is ∼ 1.4 A.U. (at the emission wavelength, i.e. 650 nm). Every photon detected by any of the 25 elements of the SPAD array generates a TTL signal that was delivered through the dedicated channel (one channel for each sensitive element of the detector) to an FPGA-based data-acquisition card (NI USB-7856R from National Instruments), which is controlled by our own data acquisition/visualization/processing software carma. The software-package carma also controls the entire bundle of microscope devices needed during the image acquisition, such as the galvanometric mirrors, the axial piezo stage, and the acoustic-optic-modulators (AOMs), and shows the data. In particular, carma synchronizes the galvanometric mirror with the photons detection to distribute photons between the different pixels of the images. All power values reported for this setup are measured at the sample plane.

To perform lifetime measurements, we also connected the five central elements of the SPAD array to a time-correlated-single-photon counting card (TCSPC) with a time resolution of 80 ps (Time Tagger 20, Swabian Instruments) operating in the so-called time-tag modality. We opportunely delayed the electronic trigger output signal provided by the excitation laser running at 80 MHz using a picosecond electronic delayer (Picosecond Delayer, Micro Photon Devices), and used the output as the reference signal (sync) for the time-resolved measurement. The carma microscope control generates the pixel, line and frame clocks, which we sent to the TCSPC card. We used a custom software module to read the stream of events outputted by the Time Tagger 20 card, each of them labeled with the corresponding inputs (sync, pixel, line, frame or element 1-5) and the time of arrival (time-tag). To reduce the data rate (and to avoid a buffer overflow), the card implemements an events filter discarding all the synchronization events (80 MHz), except those that follow one of the low rate events, thereby forming a smaller time-tag stream. Given the time-tag stream, for each photon-event its micro-time (delay from the sync signal) is calculated and the scanned lifetime histogram images (128 bins, 100 ps each), one for each SPAD element, are generated. The result is a series of three-dimensional scanned images, where the third dimension represents the lag-time (photon-arrival time) of the histogram. Relative delays between the different elements of the detector are corrected measuring their impulse-response-functions (IRFs).

**Commercial microscope**. We integrated our SPAD array into a commercial Nikon A1R confocal microscope (Supplementary Fig. S7). We did not modify the optical configuration of the system, but for our ISM detection we took advantage of the extra confocal output port of its scanning head, which has been designed for spectral detection. We removed the fiber coupling lens from the output port and we placed a motorized zoom-lens (Optem FUSION composed of a mini camera tube 3 × (f600mm), 7:1 zoom stepper motorized, and a lower lens 1 × (f200mm), Qioptiq) after the port to conjugate the pinhole plane (inside the scanning head) with the SPAD array. Thanks to the zoom lens we can add extra magnification to the system (1.3 × – 8.7 ×) in order to obtain a 1 A.U. projected size (at the emission wavelength) of the SPAD array for different objective lenses, in particular we used an CFI Apo TIRF 60 × Oil NA 1.49 (Nikon), CFI Apo LWD Lambda S 20 × WI NA 0.95 (Nikon), and a CFI Plan Apo 10 × C Glyc NA 0.5 (Nikon), in this work. The calibration of the total magnification of the system (M3) for the different settings of the zoom-lens and different objective lenses is obtained using an approach based on the estimation of the relative shift between the scanned images (Supplementary Note 2). The system performs multi-color confocal imaging thanks to a laser unit equipped with a series of continuous-wave lasers (405 nm, 488 nm, 561 nm, 640 nm). Fluorescence is filtered by the internal dichroics and bandpass filters (525/50 and 595/50 nm) before reaching the SPAD array. The control, visualization and processing are performed by the same software/hardware used for the custom system, i.e. the carma control unit. Within the Nikon application, carma switches between two modalities, namely master and slave. In the first modality, carma (i) provides analog signals to the Nikon control unit to drive the galvanometer mirrors, the stage and the AOMs, (ii) records signal from the SPAD array, (iii) visualizes and processes the data. In the slave modality, carma (i) receives the reference signal (pixel, line, and frame clocks) from the Nikon control unit which acts as a master to drive all the devices, (ii) records signal from the SPAD array, (iii) visualizes and processes the data. This latter modality has been used to implement fast ISM imaging with the resonant scanner.

**Image reconstruction**. To reconstruct the high-resolution ISM image, we implemented the pixel-reassignment method (Supplementary Note 2), which mainly consists in (i) shifting each scanned images (*i, j*) of a shift vector **s**_*i,j*_; (ii) adding up all the shifted images. The shift vectors can be estimated on the basis of the geometrical properties of the SPAD array and optical characteristic of the scanning microscope; however, in this work, we use a phase correlation approach (Supplementary Note 2) able to automatically estimate the values directly from the scanned images and able to compensate for distortions (miss-alignments and aberrations) of the system which may arise during imaging.

In the case of FLISM, the high-resolution three-dimensional image (x,y,lag-time) is obtained by applying the pixel reassignment approach to every single temporal image (x,y). The same shift vectors are used across the whole 3D stack and they are estimated on the scanned images obtained by integrating the photons across the lag-time. We finally analyzed both lifetime images with the FLIMfit software tool developed at Imperial College London (https://www.flimfit.org). Alternatively, the high-resolution ISM image can be reconstructed *via* multi-image deconvolution iterative algorithm (2–4) (Supplementary Note 4)

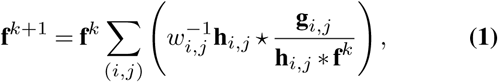

where **h**_*i,j*_ is the PSF linked to the element (*i, j*) of the SPAD array, **g**_*i,j*_ the series of scanned images, **f**^*k*^ the reconstructed image at the iteration *k*, and *w*_*i,j*_ = (0*,…,* 1] a scaling factor, that takes into account the different SNR of the images. As PSFs we used a simple Gaussian model which consider the shift vector calculated via the phase correlation method. Only few iterations of the algorithm have been applied to avoid the artifacts.

We performed the Fourier ring correlation analysis using a pair of ISM data set (series of scanned images) registered “simultaneously” thanks to a pixel dwell-time splitting (1). The fixed 1/7 threshold has been used for all the analysis.

The pixel-reassignment method, the multi-image deconvolution method, and the FRC analysis, have been implemented on the carma platform and Matlab. The Matlab source code is available upon request.

**Sample preparation**. We measured the PSF of our ISM system using gold beads and we demonstrated the enhancement in spatial resolution on two-dimensional (2D) imaging of fluorescent beads, tubulin filaments and mitochondria. We also performed three-dimensional (3D) imaging of an optically cleared mouse brain. *Gold beads.* We dropped a solution of 80 nm diameter gold beads in water onto a poly-L-lysine (Sigma) coated glass coverslip. We then mounted it with the same oil used as a medium for the objective lens (Nikon immersion oil). *Fluorescent beads.* In this study we used a commercial sample of ATTO647N fluorescent beads with a diameter of 23 nm (Gatta-BeadsR, GattaQuant). *Tubulin filaments in fixed cells.* Human HeLa cells were fixed with Ice Methanol, 20 min at −20°C and then washed 15 min for three times in a phosphate buffered saline PBS. After 1 hour in room temperature, the cells were treated in a solution of bovine serum albumin (BSA) at 3% and Triton 0,1% in PBS (blocking buffer). The cells were then incubated with the monoclonal mouse anti-*α*-tubulin antiserum (Sigma Aldrich) diluted in a blocking buffer (1:800) for 1 h at room temperature. The *α*-tubulin antibody was revealed using Abberior STAR Red goat anti-mouse (Abberior) for the custom microscope or Alexa 546 goat anti-mouse (Sigma Aldrich) for the Nikon-based microscope. The cells were rinsed three times in PBS (0.1 M, pH 7.4) for 5 min. *Dual-color tubulin filaments and mitochondria in fixed cells.* After fixation and permeabilization, as described above, BSC-1 cells from African green monkey kidney were incubated with rat anti-tubulin monoclonal antibody (YL1/2) (abcam) and rabbit anti-Tom20 (FL-145) polyclonal antibody (Santa Cruz) diluted in blocking buffer at 1:100 and 1:50, respectively for 1 hour at room temperature. The next day cells were washed with PBS three times for 5 minutes each. Secondary antibodies were applied, goat anti-rat IgG (H+L) cross-adsorbed secondary antibody, Alexa Fluor 488 (Thermo Fisher Scientific) and goat anti-rabbit IgG (H+L) cross-adsorbed secondary antibody, Alexa Fluor 568 (Thermo Fisher Scientific) at a dilution of 1:500 for 1 hour at room temperature. The cells were then rinsed with PBS (0.1 M, pH 7.4) five times, 5 min each. *Mitochondria in live cells.* To label mitochondria, Human embryonic kidney (HEK) cells were incubated with a diluted solution of MitoTracker Deep Red FM (ThermoFisher scientific) and then imaged using the Live Cell Imaging Solution (ThermoFisher scientific). *Optically cleared brain of Thy1-eYFP-H transgenic mouse*. The CLARITY method has been used to clear the mouse brain (5). In short, after perfusion, mouse brains were post-fixed in 4% PFA overnight at 4°C and then immersed in 2% hydrogel (2% acrylamide, 0.125% Bis, 4% PFA, 0.025% VA-044 initiator (w/v), in 1X PBS) for 3 days at 4°C. Samples are degassed and polymerized for 3.5 hours at 37°C. Remove samples from hydrogel and wash with 8% SDS for 1 day at 37°C. Transfer samples to fresh 8% SDS for 21 days at 37°C for de-lipidation. Then wash samples with 0.2% PBST for 3 days at 37°C. Brains were incubated in RapiClear CS (Cat#RCCS002, SunJin Lab) for 2-3 days at room-temperature for the optical clearing. The objective lens has been immersed in water for 20× imaging and in the clearing solution for 10× imaging.

## Supplementary Note 1: Single-Photon-Avalanche-Diode Array

We designed and developed a novel single-photon detector array, specifically tailored to implement image scanning microscopy. The array is composed by a square matrix of 5 × 5 single-photon avalanche diodes (SPADs) (1), having 75 *µ*m distance (pixel pitch) and 50 *µ*m side length (pixel size) with 5 *µ*m corner radius (rounded-square active-area shape, as shown in Supplementary Fig. S1a). This geometry results in a fill-factor (i.e., the ratio between photosensitive area and total detector area) of ∼ 44%, considering the external frame, otherwise ∼ 50%. The detector array is fabricated in a 0.35 *µ*m high-voltage CMOS technology, well established in the fabrication of SPADs, also allowing for the integration of on-chip readout circuitry (2). This device directly provides 25 low-jitter digital outputs (whose rising edges are synchronous to photon detections) and has the capability to selectively switch ON or OFF any single SPAD, allowing for different detection patterns (using a dedicated serial communication interface).

We chose a relative small number of pixels (i.e. 25) since our theoretical studies show that (for a fixed 1 A.U. projected-size of the detector array) a higher number of elements would provide a marginal improvement on spatial resolution (3). On the other side, a higher number of elements likely translates into more complicated data-readout architectures, precluding a fully parallel and independent operation of each pixel (thus reducing both speed and versatility of the device). To this purpose, in our SPAD array each of the 25 elements can deliver a fully-independent digital signal each time a photon is collected. Regarding the fill-factor, it is important to highlight that: (i) in the ISM application, the fluorescent photons projected on the detector array are not uniformly spread across the whole detector, but a large part of them are concentrated in the central region, thus making the overall probability that a photon reaches the active area higher than the fill-factor itself; (ii) increasing the fill-factor, by reducing the spacing between elements, will likely deteriorate performances in terms of optical crosstalk between adjacent pixels; (iii) the collection efficiency can be substantially improved by using a micro-lenses array (MLA) in front of the detector. We are currently working on the fabrication of a high fill-factor MLA directly on the SPAD array chip (4), expecting an increase of the equivalent fill-factor to above 78% (i.e., above the theoretical value predicted by using a rectangular array of circular micro-lenses).

We characterized the SPAD array in terms of photon detection efficiency (PDE), dark-count-rate (DCR), temporal response, optical cross-talk and afterpulsing probability. In Supplementary Fig. S1b, we show the measured PDE for the central pixel, as a function of wavelength and for different excess-bias voltages (*V*_*ex*_) (other pixels exhibit similar performance). The PDE decreases increasing the wavelength, ranging from about 45% at 480 nm down to 20-15% in the 600-700 nm region (both at 6 *V*_*ex*_ excess-bias). Higher PDE values could be achieved by using different fabrication technologies, as the recently demonstrated Bipolar-CMOS-DMOS technology (BCD) (5). The dark count rate has been measured at 25°C for each array element, resulting in an average DCR value around 200 counts per second (cps). The detector temporal response is shown in Supplementary Fig. S1c, for the central pixel only (similar results are obtained for all the 25 elements). It has been acquired using an external TCSPC board (SPC-630, Becker&Hickl) and with a pulsed diode laser (32 ps of FWHM, 1 MHz of repetition rate and 850 nm of wavelength, Advanced Laser System) with all the other pixel turned ON, resulting in a time jitter below 200 ps FWHM. The optical cross-talk probability between pixels is lower than 2% among closest neighbors (in the orthogonal direction). Finally, the afterpulsing probability ranges from 6.5% when enforcing a SPAD hold-off time of 50 ns, down to 1.4% with 200 ns hold-off time. Increasing the hold-off time is beneficial for the reduction of afterpulsing probability but, as a drawback, it correspondingly reduces the maximum count-rate of the detectors.

To easily take advantage of this SPAD array in the ISM microscope, a complete and standalone detection system was developed (Supplementary Fig. S1a). It is based on two stacked printed circuit boards (PCBs): the upper one hosts the detector array chip, its bias voltage generator, the hold-off time control and the serial communication interface (to individually enable/disable pixels). The lower one hosts the power supply section, a microcontroller to manage the entire system and a set of 25 low-jitter buffers, able to drive 50 Ω impedance cables for SPAD outputs. It can be directly mounted on a multi-axis positional stage for a precise and reliable optical alignment. In the context of integration of the SPAD array into an existing confocal microscope, it is important to note that the overall size of the SPAD array sensitive are is 350 × 350 *µ*m^2^, thus the magnification values requested to obtain a projected size of ∼ 1 A.U. in the sample plane are workable. For example, for imaging with the 10 × CFI Plan Apo 10 × C Glyc NA 0.5 objective lens in the green spectral range (520 nm) the size of 1 A.U. is ∼ 1.3 *µ*m, thus an extra magnification of ∼ 27 × is requested. Which in the case of the ISM implementation on Nikon A1R is obtained thanks to the extra magnification inside the scan-head (3.9 ×) and the zoom-lens system (1.3-8.7 ×).

The communication between the detection system and our scanning microscope control system (carma) is performed through 28 shielded cables: 25 of them carrying digital signals reporting the detection of a photon by each array element, the additional 3 for the serial communication interface, used to setup the detector at the beginning of each measurement.

### Supplementary Note 2: Pixel-Reassignment

In ISM the most straightforward method to recombine the scanned images into the high-resolution image is the pixel-reassignment. To understand the pixel-reassignment method it is necessary to describe the image formation process of the image scanning microscope. Since the fluorescent image scanning microscope can be considered as a linear and space-invariant system, the relation between the expected (noise-free) scanned image 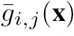 - associated to the element (*i, j*) of the detector array - and the object/specimen function *f* (**x**) can be described by a convolution operator *H*_*i,j*_

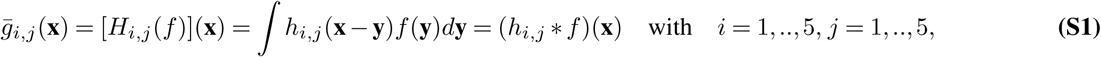

where *h*_*i,j*_ is the effective PSF associated to the element (*i, j*), **y** is the position in the sample and **x** is the position in the image back projected into the sample, i.e., the scanning position. Here, we consider a magnification equal to 1 between the object and image planes and a detector array with 5 × 5 elements. Assuming that the projected size of each element of the detector array is much smaller than 1 Airy unit (A.U.), the effective PSF of each element reads

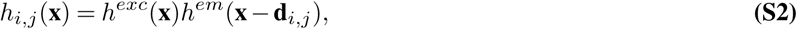

where *h*^*exc*^ and *h*^*em*^ are respectively the excitation and emission PSF of a conventional scanning microscope, and 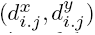 is the vector describing the displacement between the (*i, j*) element and the central element (*i* = 3*, j* = 3). For a size of the element not negligible, the emission PSF *h*^*em*^ have to be previously convoluted with the function describing the geometrical shape of the element.

For the sake of simplicity, we assume that both excitation and emission PSFs are identical, as would be the case for no fluorescent Stokes shift (i.e., excitation and emission wavelets are the same), the effective PSF will be a peak function whose maximum is located at the position midway between the excitation *h*^*exc*^(**x**) and the shifted emission *h*^*em*^(**x** -**d**_*i,j*_) PSFs’ maxima, i.e., at the position **s**_*i,j*_ = **d**_*i,j*_*/*2. Following the pixel-reassignment idea, because the signal recorded by the element i,j is most likely to have originate from the position **s**_*i,j*_, it can be “reassigned” to its original position. Performing this “reassignment” for each element corresponds to scaling the image by a factor of 2, the so-called pixel reassignment factor *α*. From the point of view of imaging, having a shifted PSF means to generate a shifted image. Thus, every scanned image is shifted, with respect the central one, by the shift vector **s**_*i,j*_ and the pixel-reassignment method can be implemented by shifting-back and adding-up the single scanned images.

This is the strategy that we implemented within this work. In particular, the shift is implemented in the Fourier domain also allowing for sub-pixel shift vectors **s**_*i,j*_. The shifting vectors can be calculated theoretically according to a simple geometrical model, i.e. the physical distance between the center of the detector elements 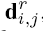, divided the magnification of the microscope on the detector plane (M3, Figure 1), and divided by the pixel-reassignment factor *α*.

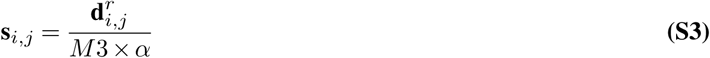

However, practically the shift vector calculated following this model significantly change from the real one. A first source of deviation is the Stokes-shift, i.e. the excitation and emission PSFs are not identical, but they have different width, meaning that the position of the maximum of the effective PSF is not located midway the excitation PSF and the shifted emission PSF and the pixel-reassignment factor is different from 2. However, a pixel-reassignment factor compensating for the Stokes-shift can be estimated *a-priori* (6). Another important source of deviation are the different aberrations which effectively change the shape of both the excitation and detection PSFs, and which are difficult to estimate *a-priori*. These aberrations influence the pixel-reassignment factor, and, more in general, the shift vectors. For these reasons, we implemented a method to estimate the shift vectors directly from the series of scanned images, **g**_*i,j*_. Clearly, this method is *a-posteriori* approach - not a real-time approach, such as for the all-optical ISM implementations, where the final image is build-up pixel-by-pixel as in a conventional confocal microscope - but it offers the important ability to compensate for system- and sample-dependent distortions. Notably, in the case of all-optical implementation based on fluorescent re-scanning two different pixel-reassigned factors can be implemented along the two-axis, but these factors need to be known a-priori.

We estimated the shift vectors **s**_*i,j*_ for the pixel reassignment using a phase correlation approach, which is typically used to estimate the drift between two images. Before describing the phase correlation approach we need to introduce the discrete notation for the scanned images. Indeed, images are usually acquired on a regular 2-dimensional raster scanning grid. If we identify each pixel by its index **n** = (*n*_*x*_*, n*_*y*_), we can denote the *N*_*x*_ × *N*_*y*_ scanned image as **g**_*i,j*_(**n**) with *n*_*x*_ = 1, …, *N*_*x*_ and *n*_*y*_ = 1*, …, N*_*y*_.

Phase correlation estimates the shift between two similar images relying on a frequency-domain representation of the data, which in our implementation is obtained through fast Fourier transform (FFT). To calculate the phase correlation between the two different scanned image **g**_*i,j*_ and **g**_3,3_, we first define the so-called correlogram **r**_*i,j*_

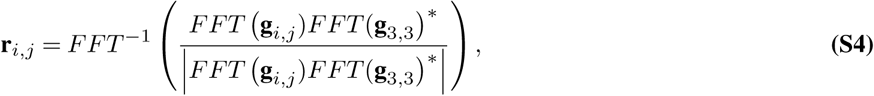

and successively find the maximum of the correlogram, whose position denotes the drift/shift between the two scanned images **g**_*i,j*_

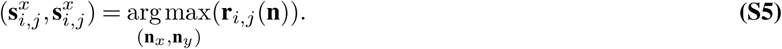

Importantly, the position of the maximum is obtained using a Gaussian-fitting algorithm or a centroid algorithm in order to obtain sub-pixel values.

Interestingly, the shift vectors can be used to estimate the magnification (M3) of the microscope on the detector array. In the absence of Stokes-shift, such as in reflection microscopy, the shift vectors are equal to half of the displacement value, **s**_*i,j*_ = **d**_*i,j*_*/*2. Since the displacement value depends on the projected physical distances of the element of the SPAD array, which are well known values, the magnification can be calculated as

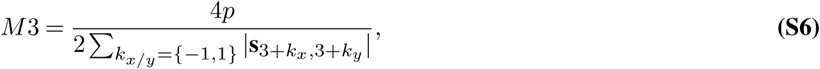

where *p* is the pixel-pitch, i.e. 75 *µ*m in our SPAD array. Only the shift vectors linked to the first-order neighbors of the central element are used, since their estimations is more robust, i.e., they are associated to higher SNR scanned images. We used reflection imaging of gold beads and this approach to calibrate the magnification for all our experiments. For example, from imaging in Figure 1b we estimate a magnification of 456×, which is fully in agreement with the set of lenses included in the custom setup (M3 = 450×).

### Supplementary Note 3: Fingerprint map

In this Note we introduce the concept of fingerprint map and we show its ability to encode information about the status of the optical scanning microscope. Such information is normally discarded in conventional scanning microscope, since single-photon detectors can register the time at which a photon reach the sensitive area but not its position. In particular, we demonstrate that from the series of scanned images it is possible to extract information about the alignment of the ISM system, and, more important, information about its PSF, regardless the specimen observed.

Given the series of scanned images **g**, we define as fingerprint map **a** the total amount of photons collected by each element of the detector array during the registration of the scanned images

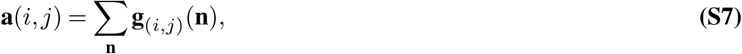

and we demonstrate that *a* is proportional to the correlation of the emission and detection PSFs of the scanning microscope. To demonstrate this proportionality we analyze the fingerprint map into the continuous domain. We consider a detector array composed by infinitesimal elements and we observe that the image *g*_*x*_′_*,y*_′ registered by the element at the position (*x*′*, y*′) *∈* ℝ^2^ reads

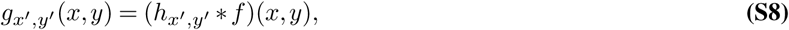

where *h*_*x′*_,_*y′*_ denotes the PSF associated with the detector element in the position (*x*′*, y*′). Thereby the fingerprint map *a*(*x*′*, y*′), defined respect to the coordinates of the detector array, reads

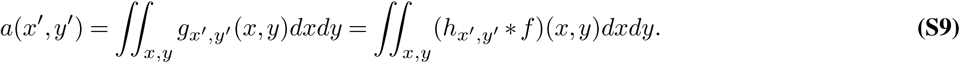

Applying the integration property of convolution, the fingerprint image reads

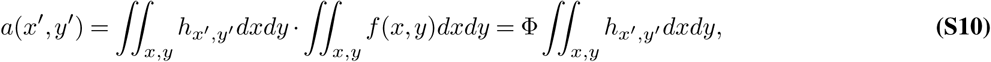

where Φ is the total flux of photons from the sample. Interestingly, *a*(*x*′*, y*′) is sample independent, at the condition Φ *>* 0, but it is strictly connected to the PSF of the microscope. Recalling that the PSF of the infinitesimal element is

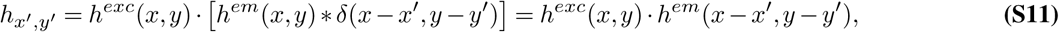

and substituting in the Equation S10, it is possible to obtain

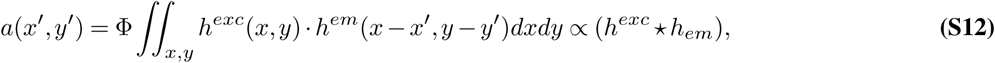

where *⋆* denotes the correlation operator. To summarize, the fingerprint image is instrument-dependent and not sample-dependent, it depends simultaneously on both the excitation and the emission PSFs.

In this work we used the fingerprint map to align the ISM system: to co-align the emission PSF with the excitation PSF we maximize the intensity of the central element (*i*=3,*j*=3) of the fingerprint map. Future directions, will be to use the fingerprint map to implement an adaptive optics (AO) feedback system, which uses the fingerprint map as figure of merit to modify light shaping devices, such as spatial-light-modulators (SLMs) and/or deformable mirrors (DMs). In a nutshell, since aberrations lead to a change of the emission and/or excitation PSFs, the fingerprint map will reflect such changes and its analysis can help in retrieving the aberrations, which can be compensated by the SLMs and/or DMs.

Another area of application of the fingerprint map is image processing, such as image deconvolution. Conventional deconvolution needs the knowledge of the microscope PSF (see 4 which is not always easy to obtain. An estimation of the PSF of the system can be decodes from the fingerprint map.

### Supplementary Note 4: Multi-image deconvolution for ISM

Another approach for recombining the scanned images into an high-resolution image is multi-image deconvolution. In comparison to pixel-reassignment, deconvolution needs higher computational effort and prior-information, such as the PSFs of the scanned images, but deconvolution can provide higher SNR and higher effective resolution (7). In this Note, we derive the multi-image deconvolution algorithm following a maximum-likelihood (statistical) approach (7) and using a discrete notation for the object function, the PSFs and the digital images.

If we denote the discretized object function, the expected scanned images and the PSFs with the vectors **f**(*n*), 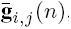, and **h**_*i,j*_(*n*), we can write the image formation process (Eq. S1) as

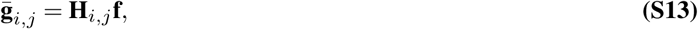

where the **H**_*i,j*_ are the convolution matrices (*N*_*x*_*N*_*y*_ × *N*_*x*_*N*_*y*_*sized*) associated with the convolution operator *H*_*i,j*_ (Eq. S1). We consider the vectors as one-dimensional vectors with *n* = *n*_*y*_*N*_*y*_ + *n*_*x*_. Moreover, discretization of convolution integral of Equation S1 using cyclic convolution and periodic extension of the pixel values of **f** and **h**_*i,j*_ reduces **H**_*i,j*_ to a circular matrix, hence the transformation

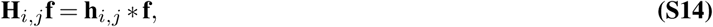

can be easily computed by means of the FFT. Here and in all subsequent equations multiplication and division of one vector by another is meant pixel-by-pixel.

The measurement process is dominated by shot noise and count rates are usually in the range of zero to a few hundred photons per pixel. Thus, for each pixel *n* and each scanned image (*i, j*), the measured value **g**_*i,j*_(*n*) is the realization of a Poisson random variable with its expectation value given by 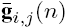. Because, each pixel is statistically independent from the other, the probability to record the series of scanned images **g** for a given specimen **f** is given by

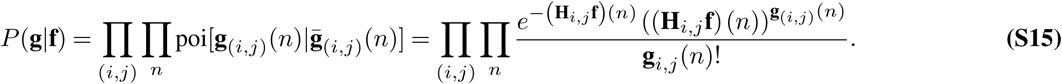

Since we assume to know the probability density *P* (**g** *|* **f**) of the data and the specimen **f** appears as a set of unknown parameters, the problem of deconvolution can be approached as a classical problem of parameter estimation, which can be solved by the standard maximum likelihood (ML) estimation approach. We introduce the likelihood function *𝔏***_g_**, defined by

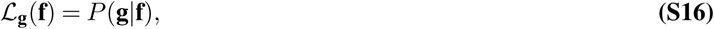

which is only a function of **f**, since the series of scanned image **g** is given. Then, the ML-estimate of the unknown object **f** is any object **f**^*\**^ that maximize the likelihood function

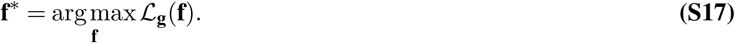

Since in our application the likelihood function is the product of a very large number of factors (Eqs. S16 and S15), it is convenient to take the logarithm of this function; moreover, if we consider the negative logarithm, the maximization problem is transformed into the minimization one. By introducing the so-called discrepancy functional *J*, the deconvolution problems reads

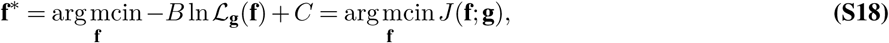

where *B* and *C* are suitable constants that can be introduced in order to simplify the expression of the functional. By using simple mathematics the discrepancy function of our application reads

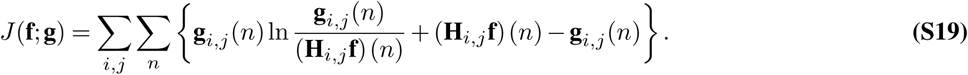

For the solution of Equation S18 we chose the split-gradient-method (SGM) (8, 9) due to its robustness, the simplicity of its implementation and its capability to enforce non-negative constraint, i.e. **f** ⩾ 0, in a natural fashion. For our discrepancy function (Eq. S19) the SGM iterations are given by

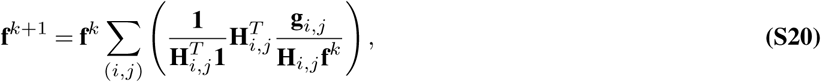

where 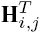 is the transpose of the operator **H**_*i,j*_ and **1** is the vector whose entries are all equal to 1. Practically, the matrix-vector multiplication 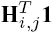 generates a vector whose elements are the sum of **H**_*i,j*_ across its columns. Since the matrix **H**_*i,j*_ is cyclic, 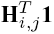 is a vector whose entries are all equal to the sum of the discretized PSF **h**_*i,j*_

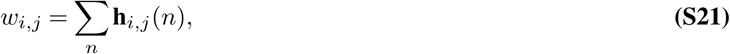

and the SGM algorithm (Eq. S20) reduces in

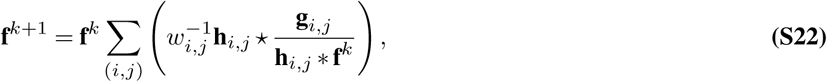

where, for the sake of simplicity, we move to a vector notation and *⋆* denotes the correlation operation, which similar to the convolution can be implemented trough FFT. The algorithm in Equation S22 can be considered as an extension of the Richardson-Lucy algorithm (10, 11) for solving the multi-image deconvolution problem. Indeed, for a single image, the algorithm reduces to the well-known RL algorithm.

Finally, it is important to discuss how we calculated the PSFs across our manuscript. We used a simplified Gaussian-based model, more rigorous model based on vectorial focusing theory (12) can be used, but they based on parameters difficult to know. For each element (*i, j*) we calculated a normalized (the integral is equal to 1) Gaussian PSF centered in **s**_*i,j*_ (Eq. S5). We used the same full-width at half-maximum for all the elements, but we scaled each PSF for a factor *w*_*i,j*_ which takes into account the expected different SNR of the associated scanned image. As scaling factors we used the values of the normalized fingerprint map, i.e. *w*_*i,j*_ = **a**(*i, j*). We estimated the FWHM directly from the images, by fitting with a Gaussian function the line intensity profiles of single isolated sub-diffraction structures in the brightest scanned image.

---

**Fig. S1.**
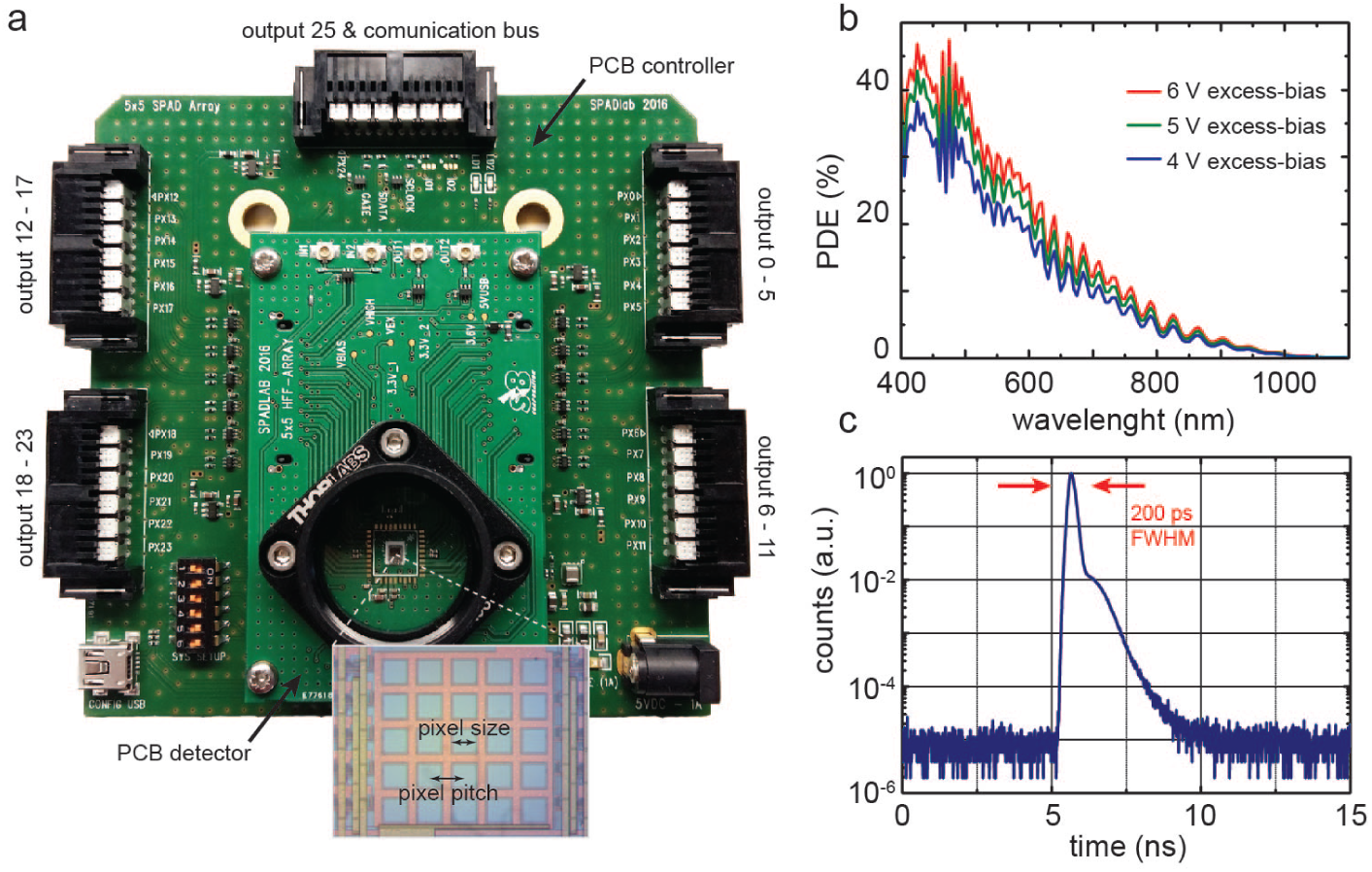
The SPAD array detection system developed for image scanning microscopy. (a) Photograph of the SPAD array detection system. The photograph shows the two stacked printed circuit boards (PCBs). Inset shows the 5×5 SPAD array. (b) Photon detection efficiency (PDE) of the central element of the 5×5 SPAD array, at different excess bias voltages. (c) Temporal response (or impulse-response-function, IRF) of the central element (*V*_*ex*_ = 6 *V* excess-bias) of the SPAD array to a pulsed laser source at 850 nm (20 ps FWHM), when all the other pixels are turned ON. The long tail on the right-side of the the IRF is due to the optical crosstalk with the other SPAD elements.

**Fig. S2.**
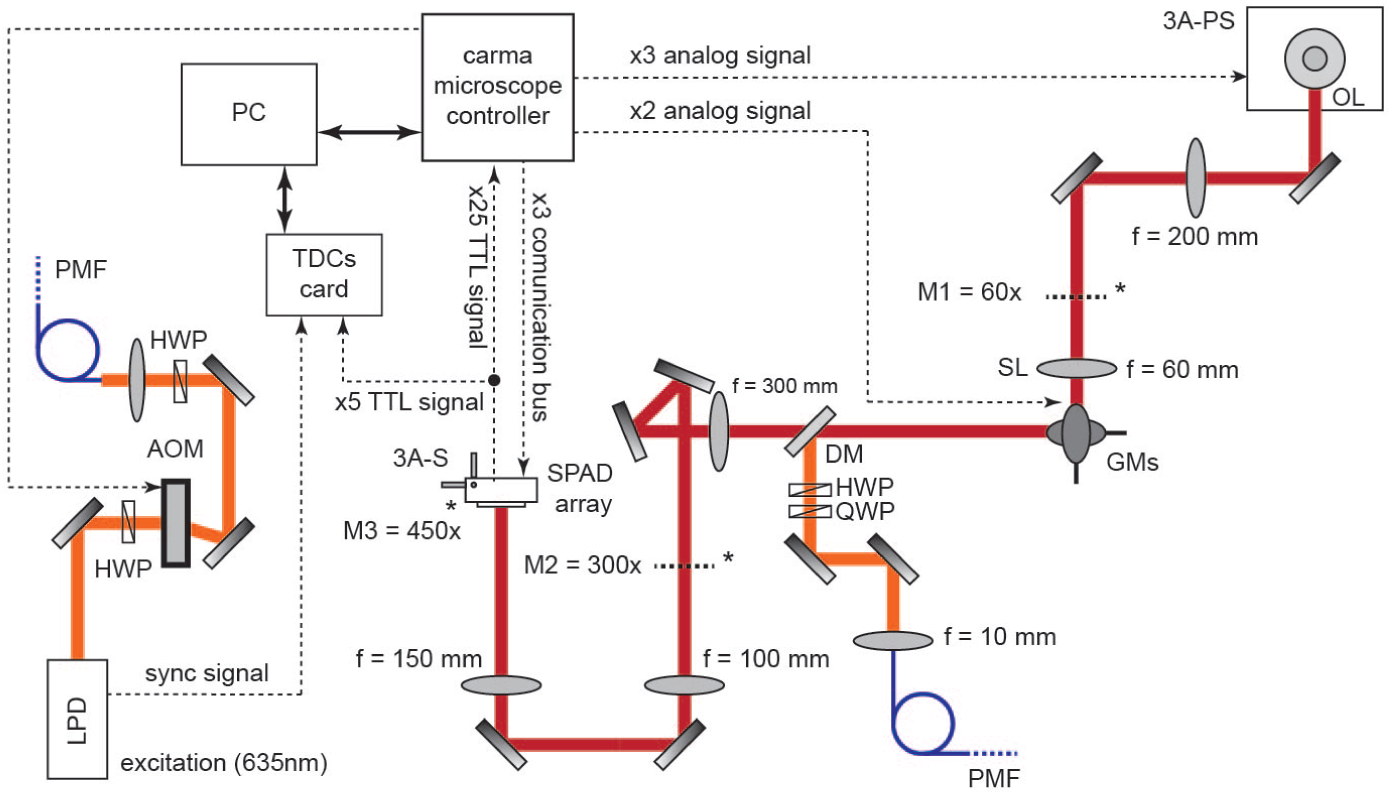
Custom image scanning microscopy schematic setup. LPD: laser pulsed diode; HWP: half-wave-plate; QWP: quarter-wave-plate; AOM: acoustic-optical-modulator; PMF: polarized maintaining fiber; PC: personal computer; TDCs: time-to-digital converters; TTL: transistor-transistor logic; 3A-S: three-axis stage; 3A-PS: three-axis piezo stage; DM: dichroic mirror; GMS: galvanometer-mirrors; SL: scan-lens; OL: objective-lens

**Fig. S3.**
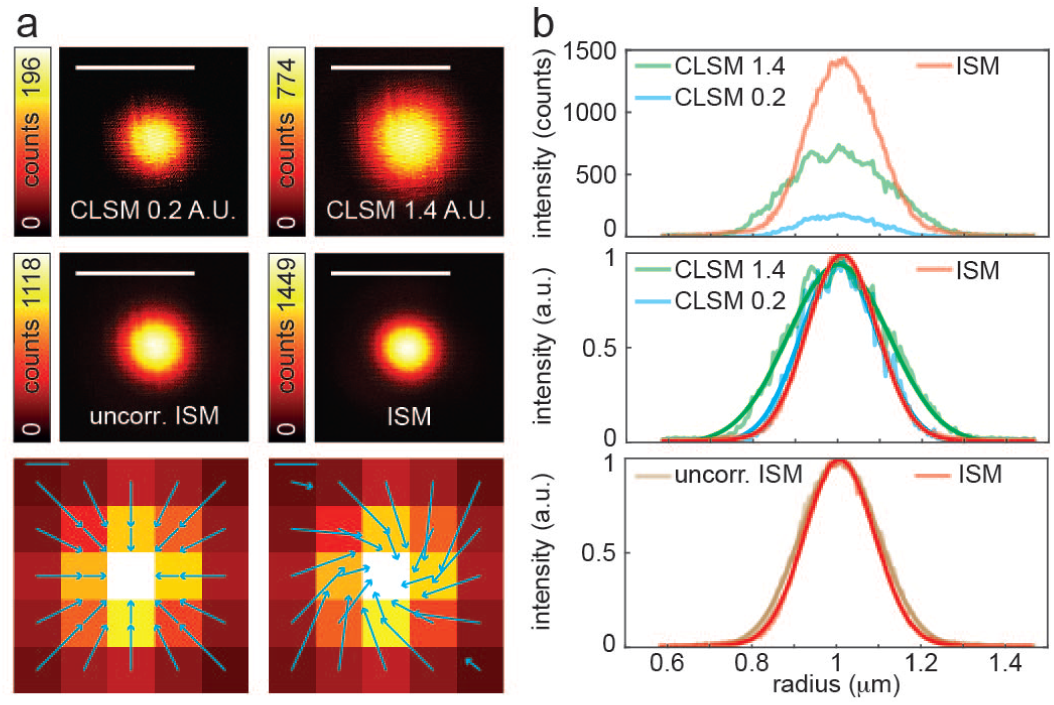
Image scanning microscopy with self-calibrating pixel reassignment. (a) Side-by-side comparison of the effective PSFs for “ideal” confocal (0.2 A.U.), “open” confocal (1.4 A.U.), uncorrected ISM (the pixel-reassignment method uses theoretical shift-vectors), and ISM (the pixel-reassignment method uses estimated shift-vectors). Scale bar: 500 nm. (b) Fingerprint maps superimposed with the shift vectors for the uncorrected ISM (left) and the true ISM (right). Scale bars: 50 nm. (c) Radial PSFs obtained from the intensity profiles of (a). The top panel shows the un-normalized PSFs, the middle and bottom panel show the normalized PSFs, together with the Gaussian fitted data. Pixel-dwell time: 50 *µ*s. Pixel-size: 5 nm. Image format: 400 × 400 pixels. Scale bars: 500 nm.

**Fig. S4.**
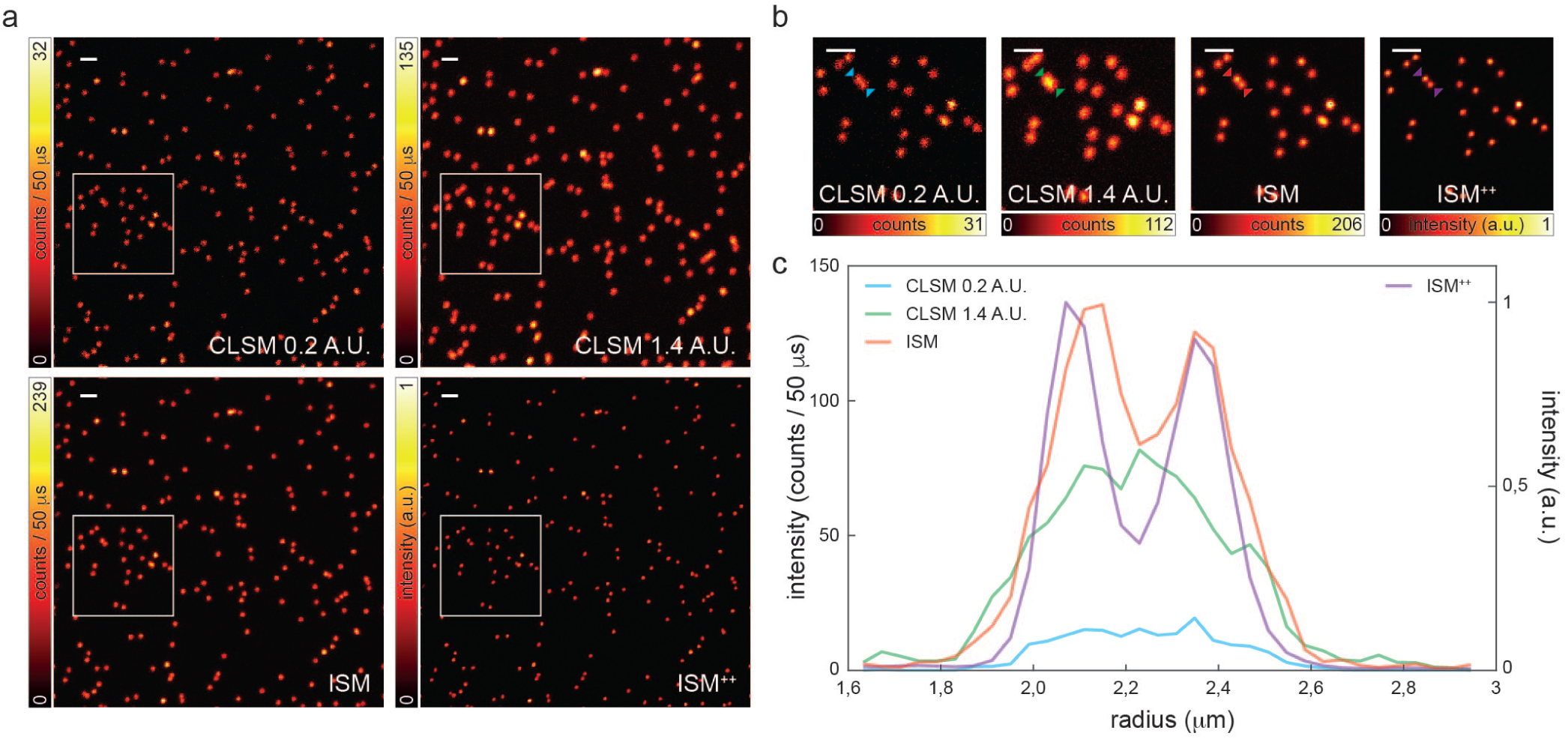
Image scanning microscopy on fluorescent beads. (a) Side-by-side comparison of “ideal” confocal, “open” confocal, ISM, and deconvolved ISM^++^ (10 iterations) images (500 × 500 pixels, 40 nm pixel size) of 20nm read fluorescent beads. Pixel-dwell time: 50 *µ*s. Pixel-size: 40 nm. Image format: 500 × 500 pixels. Excitation power *P*_*exc*_ = 56 nW. Scale bars: 1 *µ*m. (b) Magnified views of the white boxes are in (a). Scale bars:1 *µ*m. (c) Line intensity profiles across two close fluorescent beads at the position of the arrowheads in (b) for the different imaging modalities.

**Fig. S5.**
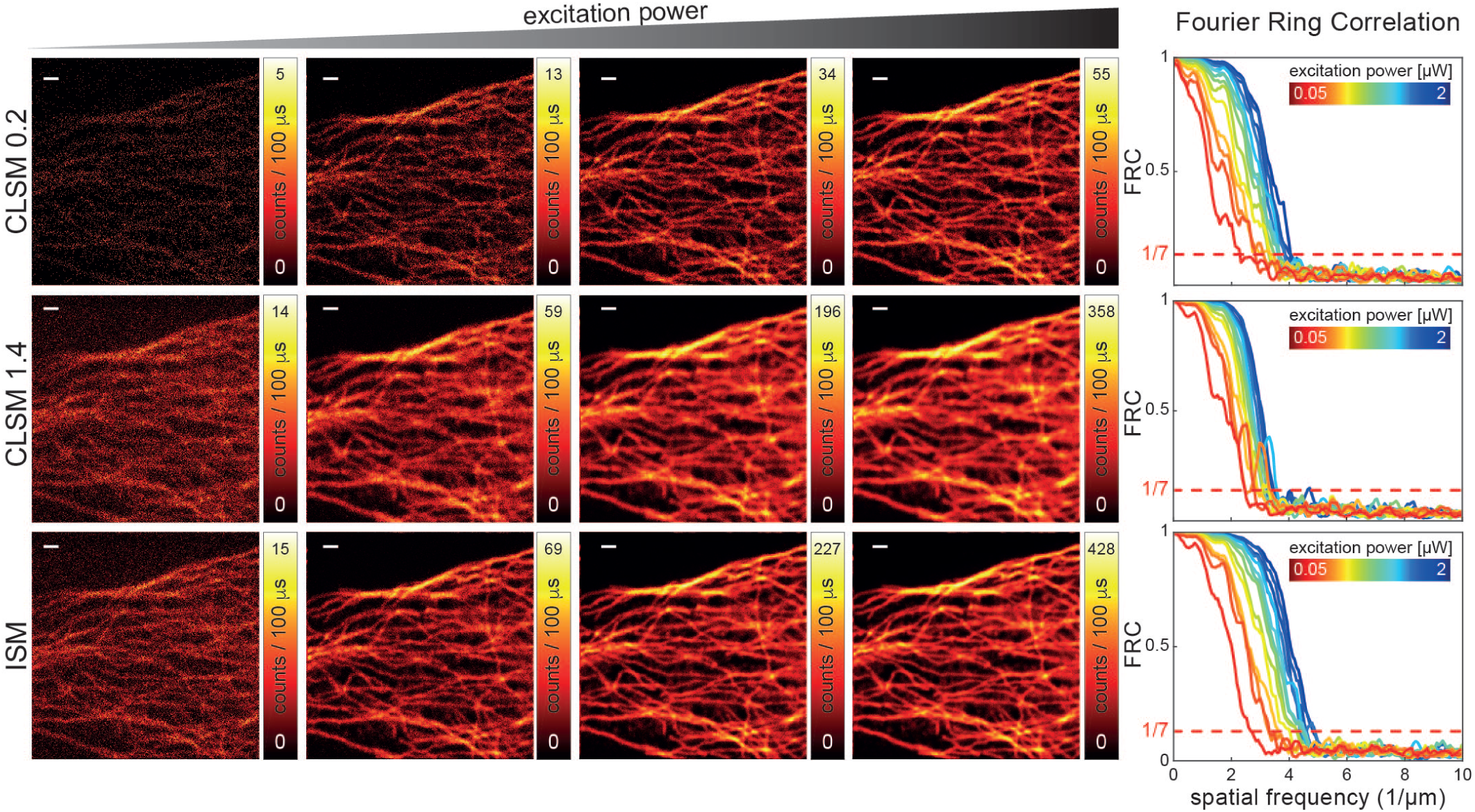
FRC-based resolution scaling on ISM with increasing excitation power. (a) Series of tubulin (labeled with Abberior STAR red) images of the same area for “ideal” confocal (top), “open” confocal (middle), and ISM (bottom) as function of the excitation beam power. Bleaching was negligible across the whole imaging experiments. Pixel-dwell time: 100 *µ*s. Pixel-size: 37,5 nm. Image format: 400 × 400 pixels. Excitation power *P*_*exc*_ = 50, 55, 62, 70, 90, 110, 170, 220, 250, 350, 520, 700, 890, 1000, 1300, 2000 nW. Scale bars: 1 *µ*m. Scale bars: 1 *µ*m. (b) Fourier-ring-correlation curves for different excitation beam powers and different imaging modalities, i.e., “ideal” confocal (top), “open” confocal (middle), and ISM (bottom). The fixed 1/7 threshold value is also reported with the curves. Resolution (based on the FRC analysis and the 1/7 threshold value) as a function of excitation power for the three different imaging modality is reported in Figure 1f.

**Fig. S6.**
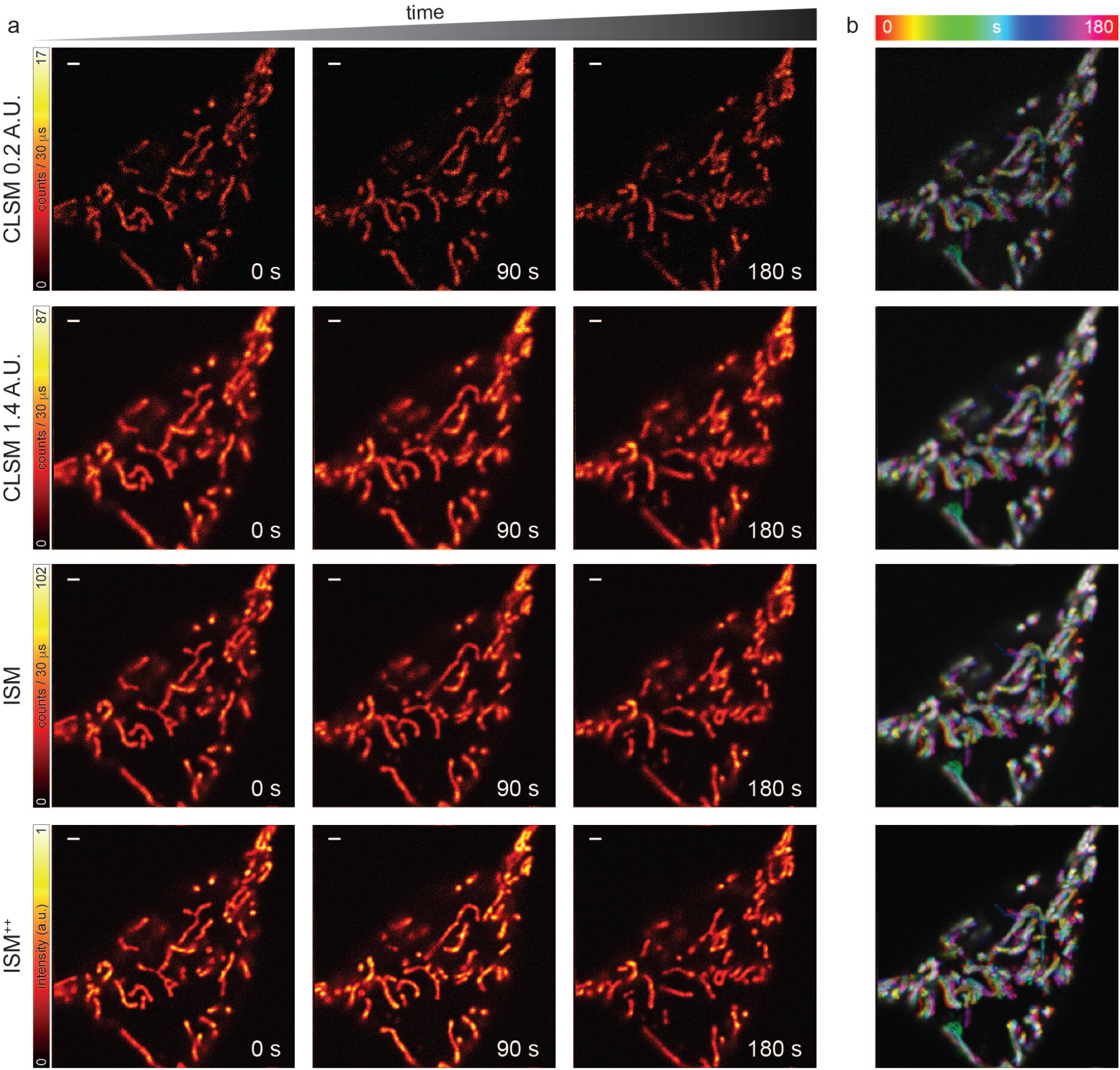
Image scanning microscopy for live cell imaging. (a) Side-by-side comparison of “ideal” confocal (top), “open” confocal (middle-top), ISM (middle-bottom), and deconvolved ISM^++^ time-lapse (3 minutes, 24 frames) of mitochondria labeled with MitoTracker Deep Red in a live-cell. For each imaging modality, three representative frames are shown (0 s, 90 s, and 180 s). Pixel-dwell time: 30 *µ*s. Pixel-size: 40 nm. Image format: 500 × 500 pixels. Excitation power *P*_*exc*_ = 140 nW. Scale bars: 1 *µ*m. (b) Maximum intensity projections (color-coded by time) of the time-lapses for the different imaging modalities. These projections allow to identify the fraction of mitochondria with minimal mobility (white) from the mobile one (colored).

**Fig. S7.**
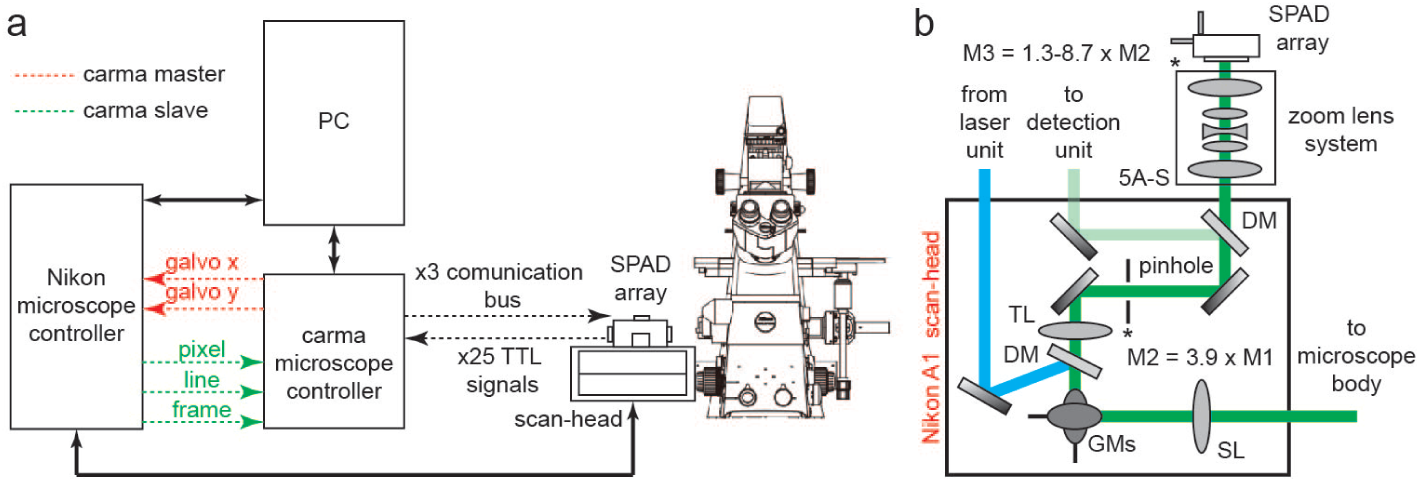
Commercial system-based image scanning microscopy schematic setup. (a) Schematic design describing the connections between the different hardware components of the image scanning microscope based on a commercial Nikon A1R system. Carma microscope control registers the photons collected by the SPAD array and it is in charge of the detector initialization. The synchronization with the scanning system and with the other actuators of the Nikon A1R, which allows generating the scanned images, is obtained through a communication with the Nikon microscope controller. In the case of imaging with galvanometer mirrors, the carma controller provides the analog scanning signal to the Nikon controller, which successively communicates with the galvanometer mirrors located into the confocal scan-head, i.e., carma is the master. In the case of imaging with a resonant mirror (for the fast axis), the carma microscope unit receives the synchronization signals (pixel, line, and frame clocks) from the Nikon microscope, i.e., carma is the slave. Both the Nikon and the carma microscope controls communicate with the personal computer (PC). In particular, the PC host the carma software, which visualizes, analyzes and processes (deconvolution, pixel-reassignment, and Fourier-ring correlation) the data. Simplified scheme of the Nikon scan-head. Only the important elements for the ISM implementation are reported. The excitation beam (blue) is sent to the galvanometer mirrors (GMs) or resonant mirror thanks to a dichroic mirror (DM). The beam is scanned on the specimen/object plane thanks to the scanning lens (SL), the tube lens and the objective lens - tube and objective lenses are not shown into the scheme). The fluorescence (green) is collected by the objective lens and de-scanned by the GMs. The SL and the TL generate a second conjugate image plane, the pinhole plane, with magnification M2 = 3.9×M1. M1 is the magnification on the first image plane (not shown in the scheme), which corresponds to the nominal magnification of the objective lens i.e, 10×, 20×, or 60× in our experiments. The DM, which usually deflects the fluorescence to the conventional single-point detector (light green), is removed and the pinhole is completely opened when performing ISM. The zoom lens (i) is positioned on a five axis-stage (5A-S) to align the fluorescence beam with respect to the SPAD array; (ii) conjugates the pinhole plane on the SPAD array and add an extra magnification (M3) on the detector plane; (iii) allows to reach a projected size of the SPAD array detector on the object plane equal to 1 Airy unit.

**Fig. S8.**
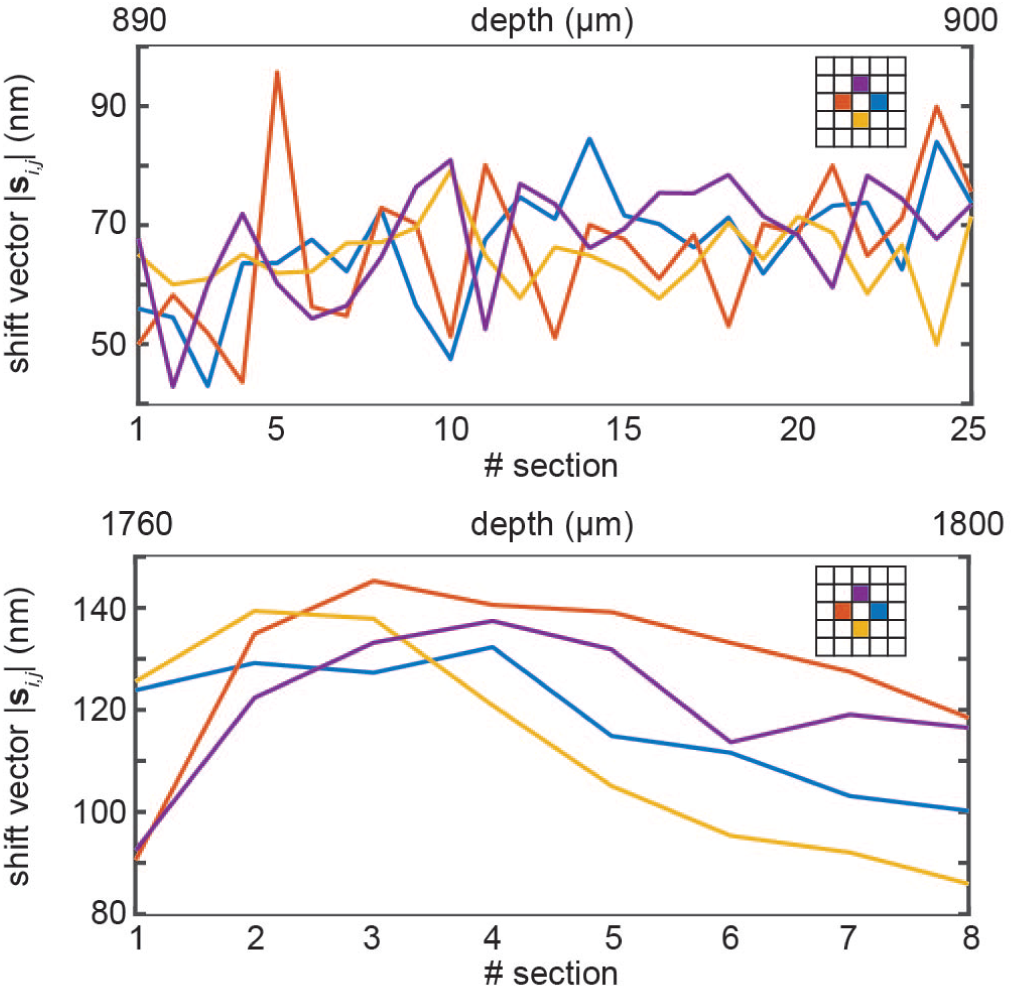
Shift-vectors as function of the depth of imaging. Module of the estimated shift-vectors for the 3D (x,y,z) dataset of Figure 3(a) (20× objective lens, top) and 3(b) (10× objective lens, bottom). The estimated shift-vectors have been calculated using the phase-correlation approach. Only the values of the four direct-neighbors element have been shown.

**Fig. S9.**
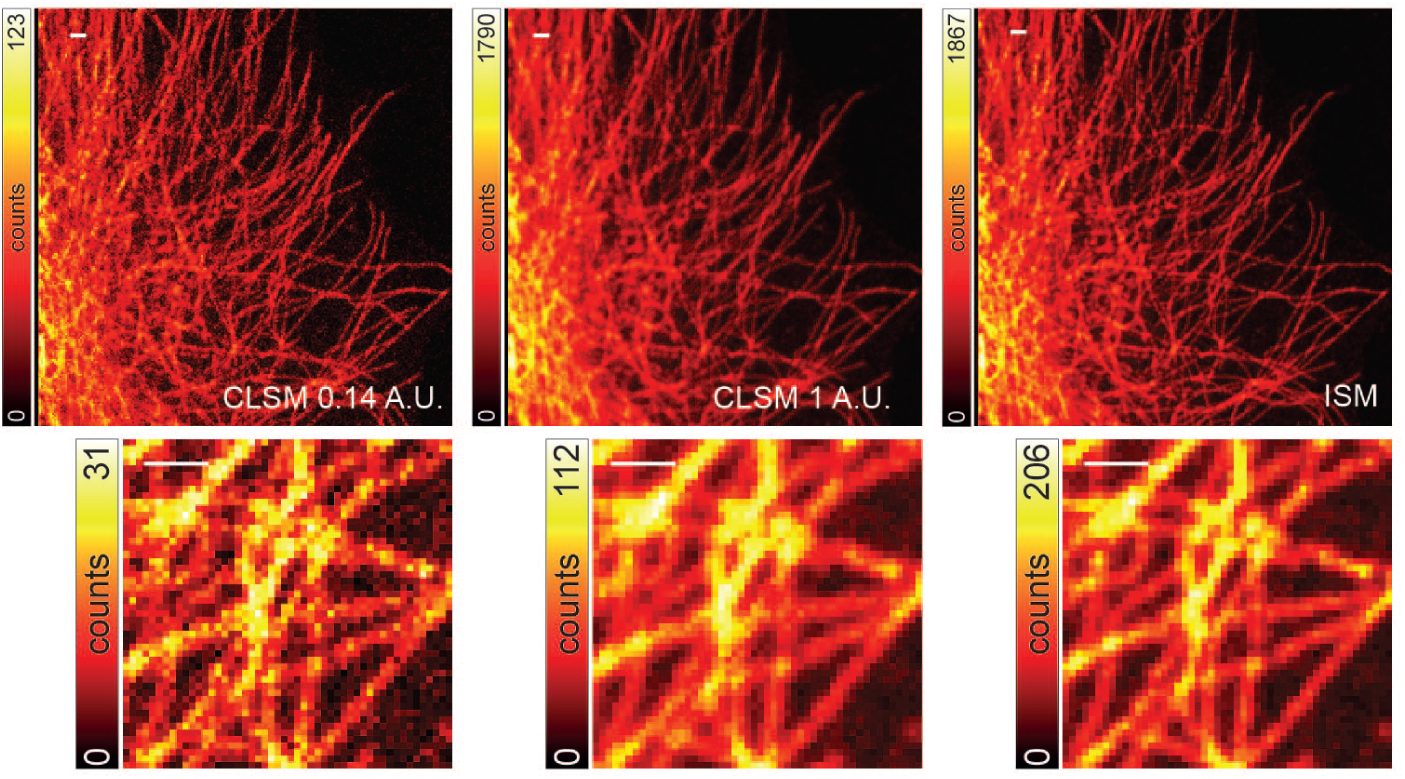
Image scanning microscopy combined with fast resonant scanning. Side-by-side comparison between “ideal” confocal, “open” confocal, and ISM images of tubulin stained with Alexa 546 (format 256×256 pixel, resonant frequency 7.9 kHz and zoom factor 8×, which results in pixel size of 103 nm and a minimum pixel-dwell time of about 70 ns, 64 line integrations). Insets show magnified views of the white boxes. Scale bars: 1*µ*m.

## Bibliography

1. Ebrecht, R., Paul, C. D. & Wouters, F. S. Fluorescence lifetime imaging microscopy in the medical sciences. Protoplasma 251, 293–305 (2014).

2. Sheppard, C. J. R. et al. Interpretation of the optical transfer function: Significance for image scanning microscopy. Opt. Express 24, 27280 (2016).

3. Sheppard, C. J. R. Super-resolutio in confocal imaging. Optik 80, 53–54 (1988).

4. Sheppard, C. J. R., Mehta, S. B. & Heintzmann, R. Superresolution by image scanning microscopy using pixel reassignment. Opt. Lett. 38, 2889–2892 (2013).

5. Müller, C. B. & Enderlein, J. Image scanning microscopy. Phys. Rev. Lett. 104, 198101 (2010).

6. Castello, M., Sheppard, C. J. R., Diaspro, A. & Vicidomini, G. Image scanning microscopy with a quadrant detector. Opt. Lett. 40, 5355 (2015).

7. Li, H., Huang, Y., Kuang, C. & Liu, X. Method of super-resolution based on array detection and maximum-likelihood estimation. Appl. Opt. 55, 9925 (2016).

8. De Luca, G. M. et al. Re-scan confocal microscopy: scanning twice for better resolution. Biomed. Opt. Express 4, 2644–2656 (2013).

9. Roth, S., Sheppard, C. J. R., Wicker, K. & Heintzmann, R. Optical photon reassignment microscopy (opra). Opt. Nanoscopy 2, 5 (2013).

10. Winter, p. W. et al. Two-photon instant structured illumination microscopy improves the depth penetration of super-resolution imaging in thick scattering samples. Optica 1, 181 (2014).

11. Gregor, I. et al. Rapid nonlinear image scanning microscopy. Nat. Methods 14, 1087–1089 (2017).

12. York, A. G. et al. Instant super-resolution imaging in live cells and embryos via analog image processing. Nat. Methods 10, 1122–1126 (2013).

13. Huff, J. The airyscan detector from ZEISS: confocal imaging with improved signal-to-noise ratio and super-resolution. Nat. Methods 12, i–ii (2015).

14. Zappa, F., Tisa, S., Tosi, A. & Cova, S. Principles and features of single-photon avalanche diode arrays. Sensors and Actuators A: Physical 140, 103–112 (2007).

15. Becker, W. (ed.) Advanced Time-Correlated Single Photon Counting Applications (Springer International Publishing, 2015).

16. Yu, Z., Liu, S., Zhu, D., Kuang, C. & Liu, X. Parallel detecting super-resolution microscopy using correlation based image restoration. Opt. Commun. 404, 139–146 (2017).

17. Sheppard, C. J. R., Castello, M., Tortarolo, G., Vicidomini, G. & Diaspro, A. Image formation in image scanning microscopy, including the case of two-photon excitation. Journal of the Optical Society of America A 34, 1339 (2017).

18. The quest for quantitative microscopy. Nature Methods 9, 627–627 (2012).

19. Tortarolo, G., Castello, M., Diaspro, A., Koho, S. & Vicidomini, G. Evaluating image resolution in sted microscopy. Optica 5, 32–35 (2018).

20. Bertero, M., Boccacci, P., Desiderá, G. & Vicidomini, G. Image deblurring with poisson data: From cells to galaxies. Inverse Probl. 25, 123006 (2009).

21. Castello, M., Diaspro, A. & Vicidomini, G. Multi-images deconvolution improves signal-to-noise ratio on gated stimulated emission depletion microscopy. Appl. Phys. Lett. 105, 234106 (2014).

22. Vicidomini, G., Bianchini, P. & Diaspro, A. STED super-resolved microscopy. Nat. Methods 15, 173–182 (2018).

23. Booth, M. J. Adaptive optical microscopy: the ongoing quest for a perfect image. Ligth Sci. Appl. 3, e165 (2014).

24. Digman, M. A. & Gratton, E. Lessons in fluctuation correlation spectroscopy. Annu. Rev. Phys. Chem. 62, 645–668 (2011).

25. Israel, Y., Tenne, R., Oron, D. & Silberberg, Y. Quantum correlation enhanced super-resolution localization microscopy enabled by a fibre bundle camera. Nat. Commun. 8, 14786 (2017).

26. Hoebe, R. A. et al. Controlled light-exposure microscopy reduces photobleaching and phototoxicity in fluorescence live-cell imaging. Nat. Biotechnol. 25, 249–253 (2007).

## Bibliography

1. Tortarolo, G., Castello, M., Diaspro, A., Koho, S. & Vicidomini, G. Evaluating image resolution in sted microscopy. Optica 5, 32–35 (2018).

2. Vicidomini, G., Schmidt, R., Egner, A., Hell, S. & Schönle, A. Automatic deconvolution in 4Pi-microscopy with variable phase. Opt. Express 18, 10154–10167 (2010).

3. Castello, M., Diaspro, A. & Vicidomini, G. Multi-images deconvolution improves signal-to-noise ratio on gated stimulated emission depletion microscopy. Appl. Phys. Lett. 105, 234106 (2014).

4. Koho, S., Deguchi, T. & Hanninen, P. E. A software tool for tomographic axial superresolution in STED microscopy. J. Microsc. 260, 208–218 (2015).

5. Chung, K. et al. Structural and molecular interrogation of intact biological systems. Nature 497, 332–337 (2013).

## Bibliography

1. Zappa, F., Tisa, S., Tosi, A. & Cova, S. Principles and features of single-photon avalanche diode arrays. Sensors and Actuators A: Physical 140, 103–112 (2007).

2. Villa, F. et al. CMOS SPADs with up to 500 µm diameter and 55% detection efficiency at 420 nm. Journal of Modern Optics 61, 102–115 (2014).

3. Castello, M., Sheppard, C. J. R., Diaspro, A. & Vicidomini, G. Image scanning microscopy with a quadrant detector. Opt. Lett. 40, 5355 (2015).

4. Surdo, S., Carzino, R., Diaspro, A. & Duocastella, M. Single-shot laser additive manufacturing of high fill-factor microlens arrays. Adv. Opt. Mater. 6, 1701190 (2018).

5. Sanzaro, M. et al. Single-photon avalanche diodes in a 0.16 µm BCD technology with sharp timing response and red-enhanced sensitivity. IEEE J. Sel. Topics Quantum Electron. 24, 1–9 (2018).

6. Roth, S., Sheppard, C. J. R., Wicker, K. & Heintzmann, R. Optical photon reassignment microscopy (opra). Opt. Nanoscopy 2, 5 (2013).

7. Bertero, M., Boccacci, P., Desiderá, G. & Vicidomini, G. Image deblurring with poisson data: From cells to galaxies. Inverse Probl. 25, 123006 (2009).

8. Vicidomini, G., Boccacci, P., Diaspro, A. & Bertero, M. Application of the split-gradient method to 3D image deconvolution in fluorescence microscopy. J. Microsc. 234, 47–61 (2009).

9. Vicidomini, G. Three-Dimensional Image Restoration inFluorescence Microscopy. Ph.D. thesis, DISI, University of Genoa (2008).

10. Richardson, W. H. Bayesian-based iterative method of image restoration. J. Opt. Soc. Am. 62, 55–59 (1972).

11. Lucy, L. B. An iterative technique for the rectification of observed distributions. Astron. J. 79, 745 (1974).

12. Leutenegger, M., Rao, R., Leitgeb, R. A. & Lasser, T. Fast focus field calculations. Opt. Express 14, 11277–11291 (2006).

